# Bumble bees (*Bombus terrestris*) use time-memory to associate reward with color and time of day

**DOI:** 10.1101/2023.04.15.536884

**Authors:** Ozlem Gonulkirmaz-Cancalar, Oded Shertzer, Guy Bloch

## Abstract

Circadian clocks regulate ecologically important complex behaviors in honey bees but it is not clear to what extent these observations can be extended to other species of bees. One key behavior is time-memory allowing foraging bees to precisely time flower visitation to periods of maximal pollen or nectar availability and reducing the costs of arriving at a flower patch at the wrong time. It is unclear whether other bees such as bumble bees, who live in smaller societies and forage over shorter distances than honey bees, can similarly associate a reward with time of day. We trained individually marked bumble bee (*Bombus terrestris*) workers to forage for sugar syrup in a flight cage with yellow or blue feeders rewarding either during the morning or evening. After a two-weeks training session, we recorded all visitations to colored feeders filled with only water. We repeated this experiment twice, with different colonies. We found that bees tended to show higher foraging activity during the morning and evening training sessions compared to other times during the day. Trained bees were more likely to visit feeders with colors rewarding compared to non-rewarding at the same time of day during the training sessions and with relatively fewer mistakes. Our findings lend credence to the hypothesis that efficient time-memory is not limited to species such as honey bees that evolved sophisticated social foraging behaviors over large distances.

## Introduction

Endogenous rhythms of about a day (circadian) are ubiquitous in plants, fungi, and animals. It is commonly accepted that these rhythms are functionally significant because they allow organisms to coordinate their physiology and behavior with the 24 hours rotation of planet earth around its axis (Benirschke, 2004; Helm et al., 2017; Sharma, 2003; Yerushalmi & Green, 2009).

Many behaviors are influenced by circadian clocks. For example, in the best-studied insect, *Drosophila melanogaster*, it has been shown that many behaviors including sleep, locomotor activity, learning, eclosion, feeding, courtship, mating, egg-laying, and time-memory are regulated by circadian pacemakers (Allada and Chung, 2010; Dubowy and Sehgal, 2017; Manjunatha et al., 2008; Saunders et al., 2002; Tataroglu and Emery, 2014). In honey bees, another important model for clock-regulated behavior (Saunders et al., 2002), the circadian clock influences complex behaviors such as sun-compass orientation, timing visits to flowers, dance communication, social division of labor among worker bees (Bloch, 2009; Moore, 2001), and queen reproductive behavior (Shpigler et al., 2022). For example, their sophisticated “waggle dance” conveys azimuth information by referring to the location of the sun and they use it to find the way to flowers 10 km away from their hive (Beekman and Ratnieks, 2000; Frisch, 1967) Here, we focus on time-memory that has been explored in diverse animal species (Biebach et al., 1989; Chouhan et al., 2015; Harrison and Breed, 1987; Reebs, 1996; van der Zee et al., 2008). It is assumed that time-memory improves foraging efficiency and thus decreases energy expenditure and predation risk and increases reward acquisition. Already at the beginning of the twentieth century, it was observed that honey bees have daily foraging rhythms (von Buttel-Reepen, 1900), and later time training experiments had established that they can associate time of day with nectar (or sugar syrup) reward at a specific location (Beling, 1929; Wahl, 1932). This complex behavior is commonly termed “time-memory”, “time-sense” or “Zeitgedächtnis” (the original name coined in German). Later studies extended this finding to show that honey bees can learn up to 9 different times of day or to time visits to different locations (reviewed in Moore, 2001). Although time-memory was first described in the Western honey bee, *Apis mellifera*, it is yet unknown to what extent this capacity is general to other species of bees.

Here we asked whether bumble bees use similar time-memory to optimize their foraging activity during the day. Bumble bees are important pollinators of agricultural crops and natural ecosystems (Woodard et al., 2015). By contrast to honey bees, their colonies are annual and are about two orders of magnitudes smaller (several hundred compared to several tens of thousands of bees per colony, respectively; Cueva Del Castillo et al., 2015). The division of labor among bumble bee workers is influenced more by body size than by age (Chole et al., 2019; Michener, 1974). Consistent with their annual life cycle and small colonies, bumble bees typically forage in their nest vicinity and rarely exceed a distance of 800 m (Walther-Hellwig and Frankl, 2000; Wolf and Moritz, 2008). Thus, the cost of mistakenly arriving at flowers at a time of low or no reward can be assumed to be lower than for long-distance foraging honey bees. There is no evidence that bumble bees can communicate the direction to floral resources in a way comparable to the honey bee dance communication (Dornhaus and Chittka, 2004, 2005; Renner and Nieh, 2008).

It is also yet unknown whether bumble bees use time-compensated sun-compass orientation to improve their foraging efficiency.

We hypothesized that *Bombus terrestris* has time-memory capacity. To test this hypothesis, we trained bumble bee foragers in a flight cage to associate yellow or blue artificial flowers with the time of a sugar syrup reward. The results of our experiments are consistent with this hypothesis.

## Materials and Methods

### Colony maintenance and setup

We conducted two experiments, each with a different colony, during September 2020 and July 2021 in the Bee Research Facility of the Hebrew University of Jerusalem, Givat Ram, Israel. Bumble bee colonies were purchased from Bio-Bee Biological Systems Ltd., Kibbutz Sde-Eliyahu, Israel. We received the colonies approximately 2–4 days after the emergence of the first worker. Each colony contained a queen, 5–10 workers, and brood at various stages of development. We transferred the colonies into wooden nest boxes (30 × 23 × 20 cm) with transparent Plexiglass covers. The colonies were provisioned with *ad libitum* pollen paste (pollen collected by honey bees mixed with commercial sugar syrup from Bio-Bee Biological Systems Ltd., Kibbutz Sde-Eliyahu, Israel). We began the training sessions when the colonies contained approximately 50 workers. We added commercial sugar syrup into 3-4 nectar pots in the colony right after the evening session in order to avoid food shortage during the night. We kept the colonies under constant darkness in an environmental chamber (27—29°C; 40—60% relative humidity [RH]). We used dim red light during feeding, inspection, or when observing the colonies. To start the training, we connected the colonies to an external flight cage with a transparent PVC tube (**Fig. 1A**).

**Figure 1.**
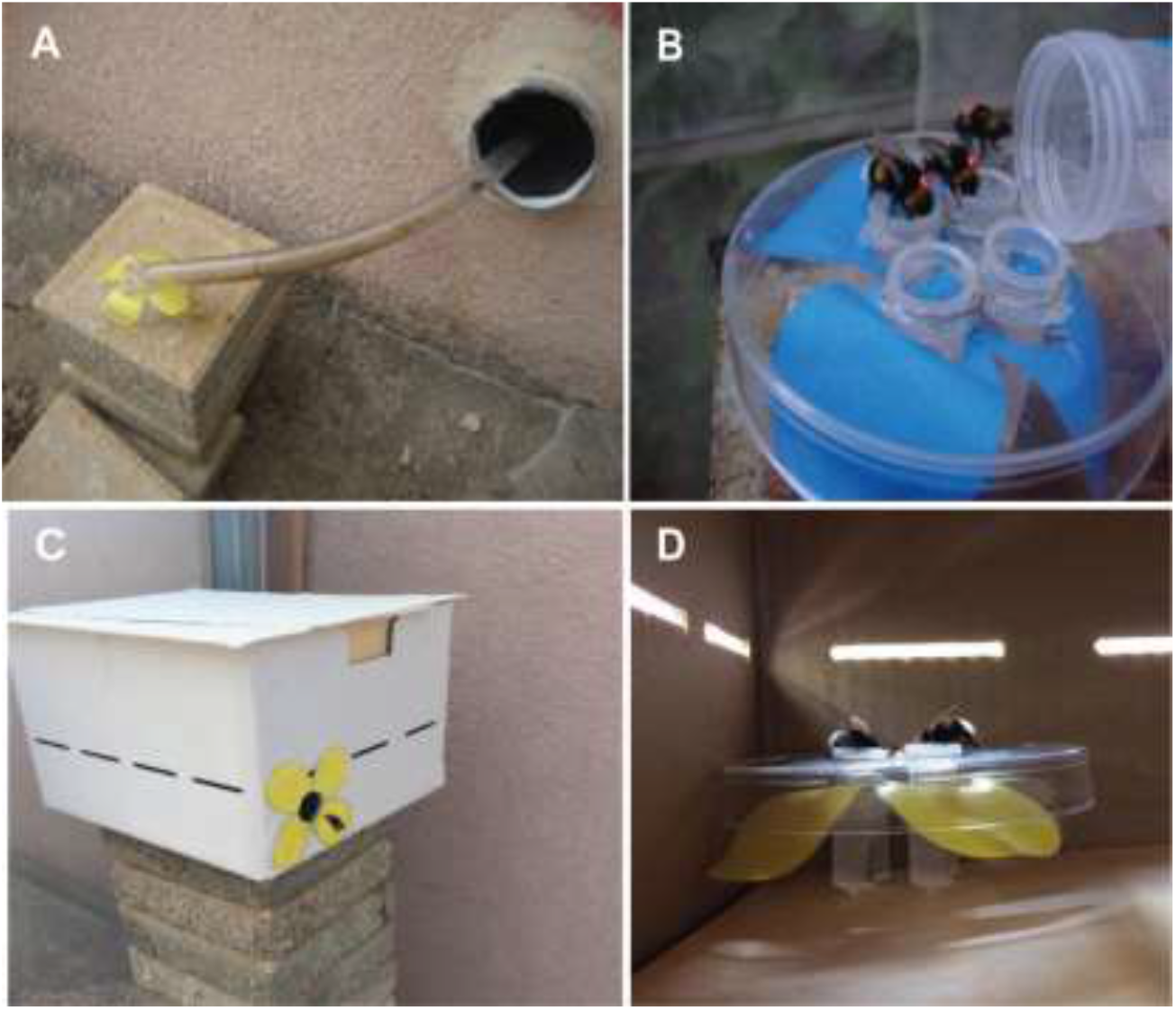
Methodological details for the first experiment testing time-memory in *Bombus terrestri*s. The tube connecting the colony to the flight cage with an artificial yellow flower at its distant end. **B)** A feeder that was filled with sugar syrup on training days and with only water on the test day (on top of a blue artificial flower). Three tagged workers are seen feeding from the feeder. **C)** A flower box in the flight cage. A trained worker is seen while flying into the opening. **D)** A look inside a flower box. Bees are seen feeding on the artificial yellow flower inside the box.

### Time learning training

#### Experiment 1

We connected two colonies of a similar age and size to the outside with transparent PVC tubes. The tube fitted to the first colony (Control) was opened outside allowing the bees to forage freely in the area around the Bee Research Facility. The second colony was connected to an external flight cage in which they forage on artificial flowers (510 × 210 × 215 cm**; Fig. 1A**). We prepared the artificial flowers in two colors, yellow or blue. We provisioned the flowers with commercial sugar syrup (from Bio-Bee Biological Systems Ltd.). We washed the flowers, including the sugar syrup feeder with soap and 70% ethanol at the end of every day to remove any scent marks that may have been deposited by the bees (Pearce et al., 2017; Saleh et al., 2007; Schmitt, 1990). The flower shape was printed on paper using a standard office printer and was attached under a Petri dish cover. The artificial flowers were made of modified Petri dishes, each equipped with four Eppendorf centrifuge tubes (2 µl) (**Fig. 1A-B-D**).

Foragers were allowed to explore the flight cage during the first five days of the training period. The sugar syrup supply inside the nest was removed at the end of the period, stimulating the bees to search for food outside. On Day 6 (start of training), we presented artificial flowers filled with commercial sugar syrup. We first placed the flowers right at the nest entrance (**Fig. 1B**) and confirmed that foragers drink syrup from the artificial flower. Starting on Day 7, we gradually (∼every hour) moved the artificial flowers farther and further away from the nest entrance, until they reached two boxes placed in the far-off corners of the flight cage. The flowers were first placed in front of the boxes, not inside. On Day 8, we presented a rewarding yellow flower next to one of the boxes during the morning hours (7:00-9:00) and a rewarding blue flower next to the other feeder during the evening hours (16:30-18:30). The time of sunrise and sunset during the entire training period were 06:18-06:23 and 18:55-18:43, respectively (*Time and date*, 2020a). The temperature range was between 25-32°C, and the humidity range was between 52-67% (*Time and date*, 2020b) on the test day. On Day 10, we moved the flowers into the boxes (**Fig. 1C-D**). First, we left the box lids open for the first hour of the reward for each session. After observing five bees feeding inside the boxes, we narrowed down the time of food availability to one hour during both the morning and the evening training sessions and left the boxes closed after these times.

The whole procedure of tracking and training the colonies took two weeks (Supplementary **Fig. S1**). We conducted the test day at the end of the second week. During the test day, the flowers were filled with only water.

The activity of the control colony was assessed qualitatively by observations.

#### Experiment 2

The overall experimental design (Supplementary **Fig. S2**) was similar to the first experiment, with the following modifications:

- **Flower shape:** We slightly modified the artificial flowers in order to increase the number of bees that can be trained at once. We made the flowers in four petal shapes from a Petri dish and covered these plates with yellow and blue stickers instead of attaching a flower-shaped paper under a Petri dish cover as in the first experiment (**Fig. 2A-B**). Using stickers also helped reduce light reflection which was occasionally a problem with the Petri dish cover. We replaced the stickers every 2-3 days. The syrup container was a cap of an Eppendorf centrifuge tube (1.5 ul) that was glued to the plate and was filled with up to 300 µl commercial sugar syrup.
- **Acclimation:** The cage-adjustment period lasted two days. At the end of this period, we removed the sugar syrup feeder from the nest.
- **Flower presentation:** On Day 3, we started to present flowers filled with commercial sugar syrup next to the entrance tube. After that, we hung the artificial flowers by using plastic cable ties from the ceiling of the flight cage. We attracted the first few bees by allowing them to feed on a syrup-coated Pasteur pipette in order to facilitate training. We slowly transferred them from the entrance to the hanging flowers on the other side of the cage, while they were feeding on the Pasteur pipette. We removed the pipette gently right after they switch to feed on the flower. This strategy facilitated moving the trained bees from the hive entrance toward the artificial flowers. Typically, additional foragers followed the already-trained bees and landed directly on the same flower.

**Figure 2.**
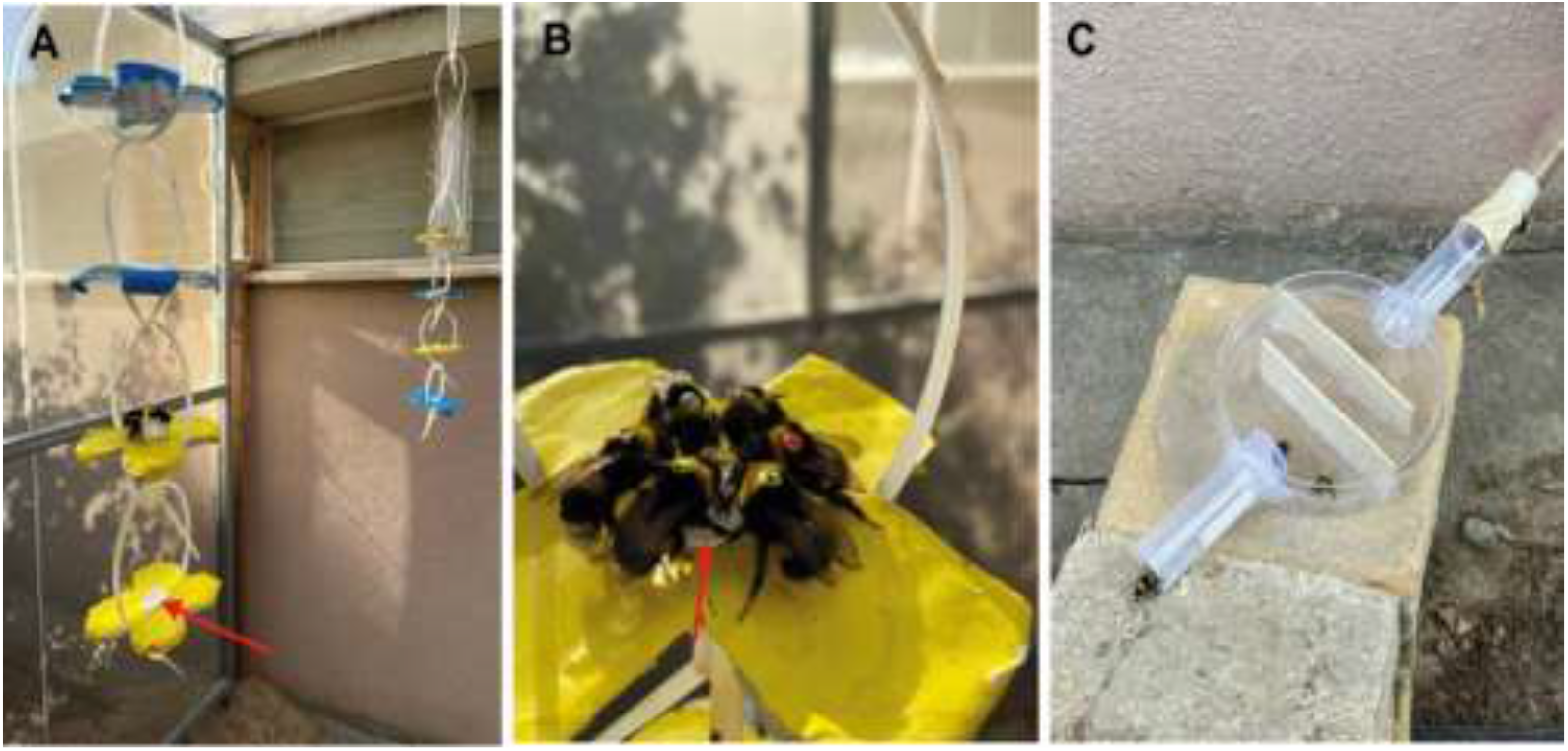
Methodological details of the second experiment. **A)** Artificial flowers painted yellow or blue and hung from the ceiling of the flight cage. Each artificial flower is equipped with a small feeder made of an Eppendorf tube cup (red arrow) in the middle. **B)** Tagged bees consuming sugar syrup on a yellow artificial flower. **C)** The maze on the entrance tube, was used to slow down bee movement on their way in or out of the nest box.

On Day 5, in which we recorded five bees that are trained to feed on the artificial flowers, we removed the yellow and the blue artificial flowers to the different corners of the flight cage (**Fig. 2A**). We tagged the workers with green numbered tags (**Fig. 2B**). On Day 8, we narrowed the period in which the flowers are rewarding to two hours in the morning (7:00-9:00) and two hours in the evening (16:30-18:30). On Day 11, we further narrowed the time during which the flowers are rewarding to one hour in the yellow flower during the morning (7:30-8:30) and one hour in the blue flower during the evening (17:00-18:00). At the same day, we also increased the number of flowers to four of each color such that more bees could obtain a reward. We changed the position of the flower colors randomly over the day to make sure the bees learn to associate the reward with the flower’s color rather than with its position in the flight cage. On Day 15, we installed a maze at the end of the entrance tube in order to slow the bees exiting and entering the nest and facilitate recording the tag details (**Fig. 2C**). One week before the test day, we tagged all recently emerged untagged bees with orange color number tags. Thus, worker bees were tagged with tags of two different colors corresponding to their age cohort. On the test day, green-tagged bees were about one week older than the orange-tagged bees. The times of sunrise and sunset during the entire training period were between 05:37-05:47 and 19:48-19:43, respectively (*Time and date*, 2021a).

On the test day, we filled the feeders with tap water (300 µl) on each flower and refilled them at the beginning of each session. We recorded all visitations to the artificial flowers during the following sessions: morning (6:30-9:30), mid-day (11:00-11:30, 13:00-13:30, 15:00-15:30), and evening (16:30-19:30). We recorded the first choice for each bee in each 30-minute intervals during each observation session, as well as all the following visits to the artificial flowers. A visit was defined as a bee landing (all legs on the flower) and her proboscis is extended out. Hovering above the flower was not counted as a visit. We also counted how many bees are going out through the tube per 30 min during these observation sessions in the focal and the control colonies.

Three observers recorded bee activity during the observation sessions on the test day. Two of them recorded visits to flowers (four flowers each) and the third one recorded the number of bees departing from the free-foraging control colony. We automatically video recorded the number of bees exiting the focal colony (connected to the flight cage) using an iPhone 11 camera which we placed on top of the maze at the end of the entrance tube. Foraging duration in the flight cage was calculated as the difference between the time of departure and the time of return for each trip of a tagged bee.

At the end of the experiment, all worker bees (except the very small-sized) in the colony were tagged (total number of tagged=109, green-tagged=86, orange-tagged=23, untagged=43) with distinct colored number tags on the dorsal part of their thorax. The temperature ranged between 26-33°C, and humidity between 44-65% during the test day (*Time and date*, 2021b). We froze (−20°C) the entire colony on the day following the test day. We measured the length of the marginal cell of the forewings (one measurement per bee) as an index for body size (as in Yerushalmi et al., 2006).

#### Learning Score

We calculated for each bee a learning score (LS) as an index for learning performance. The LS presents the proportion of correct choices and was calculated using the following equation:

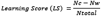

**Nc:** Number of visits to the rewarding color (correct choice)

**Nw:** Number of visits to the non-rewarding color (wrong choice)

**Ntotal:** Total number of visits in the session (Ntotal = Nc + Nw).

The score varies between 1 = only correct choices, to -1 = only wrong choices. A learning score of 0 indicates equal number of correct and incorrect choices. We focused on four time intervals during the test day: “Pre-morning” – the observations before the time of the morning training (6:30-7:30), “Morning” – the time of the morning training (7:30-8:30), “Pre-evening” – the observations before the time of the evening training (16:30-17:30), “Evening” – the time of the evening training (17:30-18:30).

### Statistical analyses

We used Pearson’s Chi-Square test to test if the frequency of visits to and the first choice of the yellow or blue flowers per observation session differ from 50%. We applied the Bonferroni correction for multiple tests.

We tested if the learning score of each time session differ from zero using one-sample Wilcoxon signed rank tests. We used the Shapiro-Wilk normality test or Kolmogorov-Smirnov normality test to determine if the data distribution is normal. The frequency distribution of bee activity at the colony entrance of the two colonies in Experiment 2 was compared using a two-sample Kolmogorov-Smirnov test. These analyses were performed in R v. 4.0.320. We used 2-way ANOVA to compare the learning score and foraging duration of the two age groups in Exp. 2. Two-way ANOVA and all the graphs were performed in GraphPad Prism version 5.0.

## Results

### Experiment 1

A total of six bees explored the cage during the test day. We limited our analyses to four of them that were recorded visiting rewarding flowers during at least two training days. We use “training session” to refer to the time of reward during the training period. The trained bees started visiting flowers at around 06:30 which is earlier than the time of the training session (**Fig. 3A**). Visitation rates increased until around 08:00, and the number of visits typically decreased towards 09:00 during which no bee was recorded in the flight cage. Only a single focal bee flew in the cage during the midday observations. The foraging activity started to increase again at around 16:00, just before the evening training session, and reached its peak between 16:30 to 17:30. The last bee returned to the hive at around 18:30. Qualitative assessment of flight out of the control colony reveal relatively high activity also during midday hours during which there was almost no activity in the flight cage. This observation suggests that time training influence the organization of foraging activity in the flight cage.

**Figure 3.**
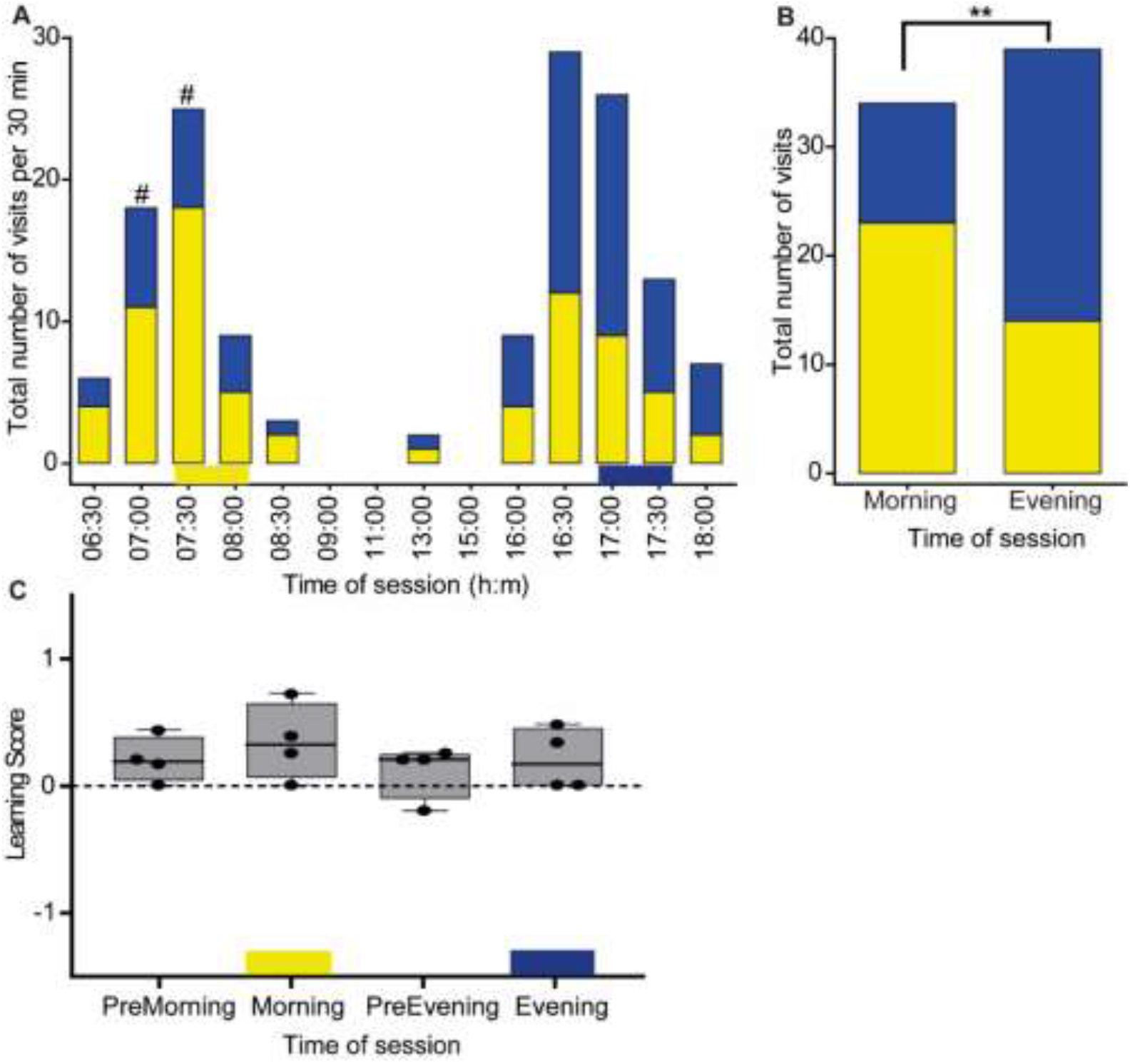
Time-color learning in Experiment 1. **A)** The total number of visits of trained bees (n=4) to the yellow or blue flower boxes as a function of time of day. The values depict the number of visits in each half-hour interval. The yellow and blue horizontal bars at the bottom depict the rewarding period and color during the training session. “#” – The probability to visit the yellow and blue flowers differed in a χ^2^ test (*p* ≤ 0.05), but not after Bonferroni corrections. **B)** A complementary analysis for pooled samples of all visitation during the times in which a reward was provided during the training (7:30-8:30 and 17:00-18:00). Asterisks denote statistical significance for Pearson’s Chi-squared independence test, \*\**p*-value = 0.006. **C)** Learning Score.

The bees visited the yellow flower more often during the time corresponding to the morning training session (visits to yellow=23, blue=11) and the blue flower more often during the time corresponding to the evening training session (visits to blue=34, yellow=21). The frequency of visiting the two colors differs from 50%:50% only for the morning session in a Chi-squared test (*p*<0.05), but this difference was not significant after Bonferroni correction for multiple comparisons (**Fig. 3A, Table S1**). When we limited our analyses to the time of training sessions, we found that the likelihood to visit a yellow or blue flower differ between the morning and evening training sessions (Pearson’s Chi-squared independence test, χ^2^ = 7.3255, df = 1, *p*-value = 0.006, **Fig. 3B**). In these analyses too, there were more visits to the yellow flower during the morning training session (Chi-squared test, *p*-value = 0.039), however, visits to the blue flower were not different than 50%:50% during the evening training session (Chi-squared test, *p*-value = 0.078). The results from the Learning Score (LS, see Materials and Methods) were positive for all the sessions, but did not differ from zero for any of the sessions (Wilcoxon signed-rank exact test: *p*-value > 0.05 for all four time-points, **Fig. 3C**).

The line in the boxplot shows the median and the box frame spans over the first to the third quartile. Each dot shows the value of a single bee. A gray shaded box indicates that the Learning Score is not statistically different from zero (one-sample Wilcoxon signed rank test, *p* > 0.05). Results are shown for visits during one-hour intervals, before the morning training (PreMorning), during the morning training (Morning), and before the evening training (PreEvening), during the evening training (Evening).

#### Experiment 2

Seventeen bees were successfully trained during the training period. Bees started departing from both the focal and control colonies early in the morning of the test day (departures at 6:30 were not recorded for the focal colony, but we can still infer their activity from the color choice plots below). Foragers of the control colony which were freely foraging were active throughout the day with a 40% drop around noon (11:00-15:30). The focal colony which was limited to forage in the flight cage, showed a stronger drop in activity level during these hours and was specifically active at and around the times of sugar syrup reward during the training session, but the difference in the frequency distributions of activity over time was not statistically significant at the p<0.05 level (Two-sample Kolmogorov-Smirnov test, D = 0.5, *p*-value = 0.06) (**Fig. 4**).

**Figure 4.**
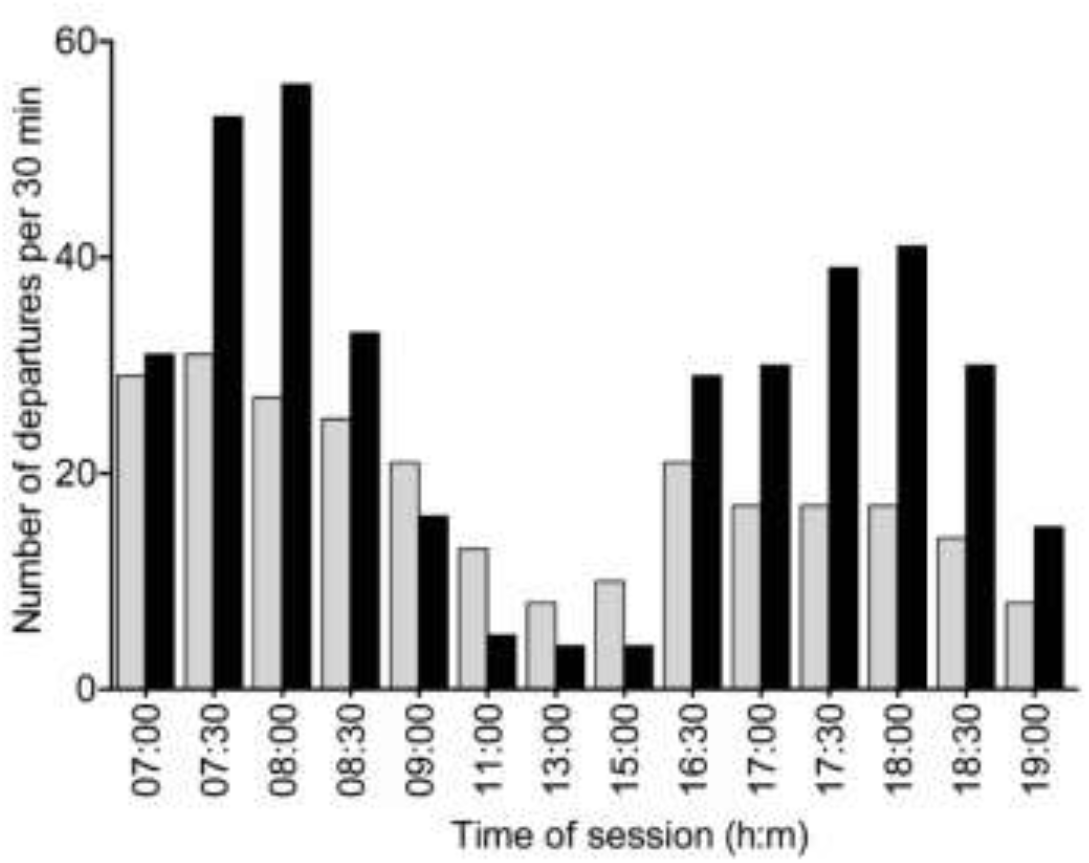
Departures over time during the test day in Experiment 2. The number of departures was recorded at the entrance tubes of the trained colony that was limited to forage in the flight cage (black bars) and the free foraging control colony (gray bars). The frequency distribution of activity did not differ between the two colonies, but shows a clear trend (Two-sample Kolmogorov-Smirnov test; D = 0.5, *p* = 0.06).

The trained bees were more likely to first visit the correct color during the four 30 minutes intervals corresponding to time of training, and the probability of selecting the correct color was statistically different from random in 3 of the 4 intervals (marked with asterisks in **Fig. 5A**). The preference for yellow was apparent already one hour before the time of the morning training (6:30-7:00 – 100%, 7:00-7:30 – 71.4%, 7:30-8:00 – 100%, 8:30-9:00 – 70%). After the morning training, the bees changed their preference and their first visit was to the blue flowers (8:30-9:00 – 82.3%, 9:00-9:30 – 62.5%). During the evening session, the choice of the blue flower was clearer (16:30-17:00 – 83.3%, 17:00-17:30 – 87.5%, 17:30-18:00 – 93.3%, 18:00-18:30 – 100%, 18:30-19:00 – 100%, 19:00-19:30 – 40%). The *p* values for each hour are calculated and shown in **Table S2**. When we limited our analyses to time of training sessions, we found that the likelihood to visit a yellow or blue flower differ between the morning and evening training sessions (Pearson’s Chi-squared independence test, χ^2^ = 110.3, df = 1, *p* < 2.2e-16, **Fig. 5C**). The total visits to the yellow flower during the morning training session differed from 50%:50% significantly (Chi-squared test, *p*-value < 2.2e-08), and visits to the blue flower during the evening training session was also significantly different than 50%:50% (Chi-squared test, *p*-value < 3.0e-19).

**Figure 5.**
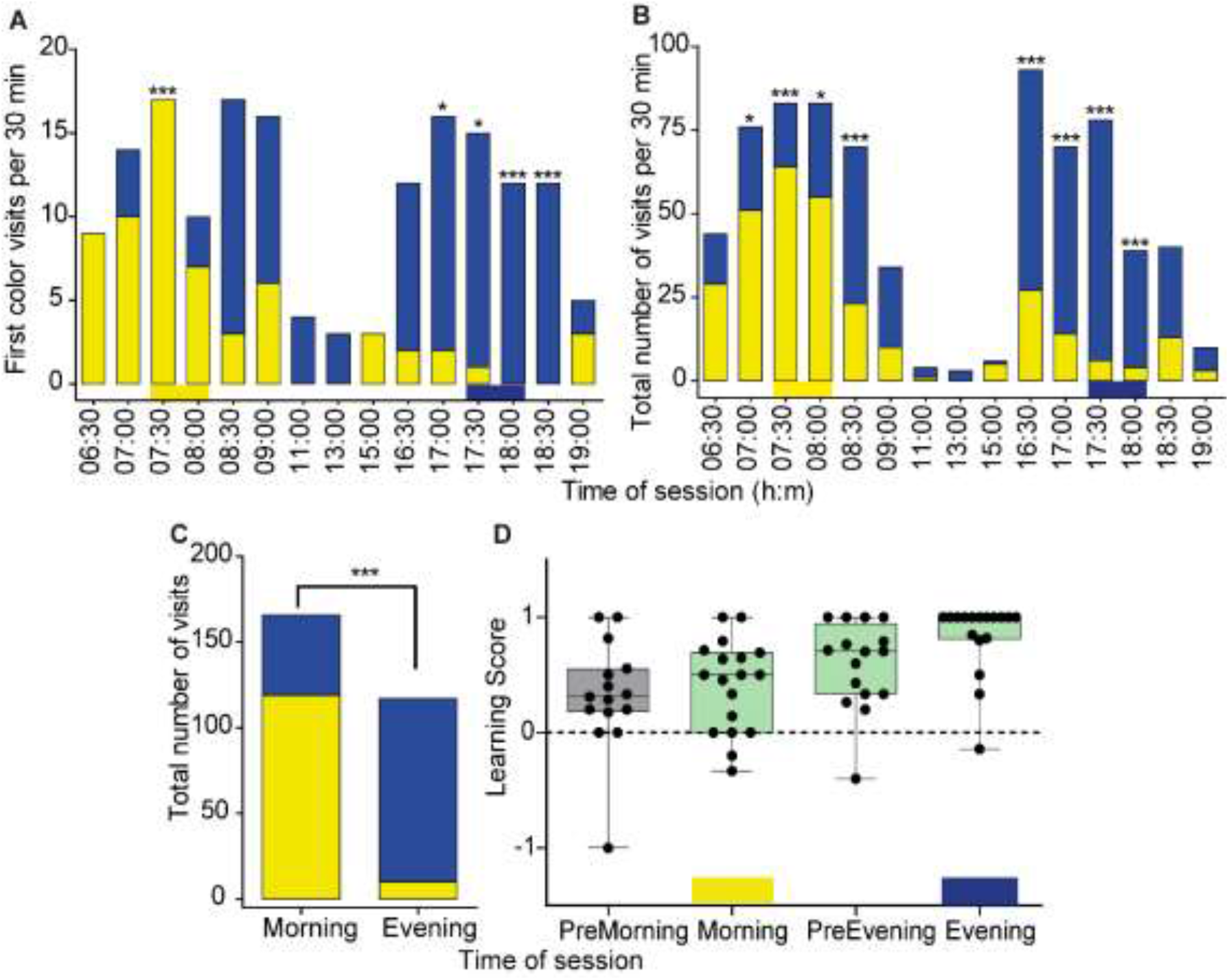
Time-color learning in Experiment 2. **A)** The first color visited by 17 trained bees. The choice was recorded for each 30 minutes observation session. The yellow and blue horizontal bars at the bottom depict the rewarding period and color during the training session. **B)** Total number of visits to the yellow and blue colors. Asterisks in A and B denote the probability to visit the yellow and blue flowers differed in a χ^2^ test: \**p* ≤ 0.05, \*\*\**p* ≤ 0.001. **C**) A complementary analysis for pooled samples of all visitation during the times in which a reward was provided during the training (7:30-8:30 and 17:30-18:30). Pearson’s Chi-Squared independence test, \*\*\**p* ≤ 0.001. **D)** Learning Score (LS) during the test day. A green box indicates that the difference between the LS and 0 is statistically significant (one-sample Wilcoxon signed rank test, *p* <0.05). Other details are as in **Fig. 3C**.

A preference for the rewarding color during the training period was also apparent when looking at the total number of visits to the yellow and blue flowers during the test day. The trained bees visited the yellow more often than the blue flowers during the morning sessions and the hour preceding it (PreMorning, 6:30-7:00 – 65.9%, 7:00-7:30 – 67.1%; Morning, 7:30-8:00 – 77.1%, 8:00-8:30 – 66.3%; **Fig. 5B**). After the morning training session, the bees preferred to visit the blue over the flowers (8:30-9:00 – 67.1%, 9:00-9:30 – 70.6%). The trained bees showed a clear preference for the blue color during the times corresponding to the evening training session (PreEvening, 16:30-17:00 – 71%, 17:00-17:30 – 80%; Evening, 17:30-18:00 – 92.3%, 18:00-18:30 – 89.7%; PostEvening, 18:30-19:00 – 87.5%, 19:00-19:30 – 70%).

The Learning Score of the trained bees was positive and different from zero during both the morning (Morning, median = 0.5, *p* = 0.02) and evening (Evening, median = 1.0, *p* = 0.002) training sessions **(Fig. 5D)**. The finding is also consistent with anticipation given that the LS was positive already during the one hour before the time of reward, although the LS was significantly different from 0 only for the period before the evening training session (PreMorning, median learning score = 0.31, Wilcoxon signed rank test with Bonferroni correction: *p* = 0.17; PreEvening, median = 0.7, *p* = 0.01).

Given that in this experiment we trained bees of two age cohorts, we further tested if age or foraging experience influences learning performance. The training data of the two cohorts are summarized in **Table S3**. We found that both age cohort and time of day affected the foraging duration of the trained bees with a significant interaction between these two factors (Two-way ANOVA; age, *p* = 0.01; time of day, *p* = 0.01; interaction, *p* = 0.02; **Fig. 6A**). The interaction suggests that the influence of time of day on foraging duration was different for the two age cohorts. There was no effect of the age cohort on learning performance in a test in which we focused only on the times of the morning and evening training sessions (Unpaired t-test, *p* = 0.16, **Fig. 6B**).

**Figure 6.**
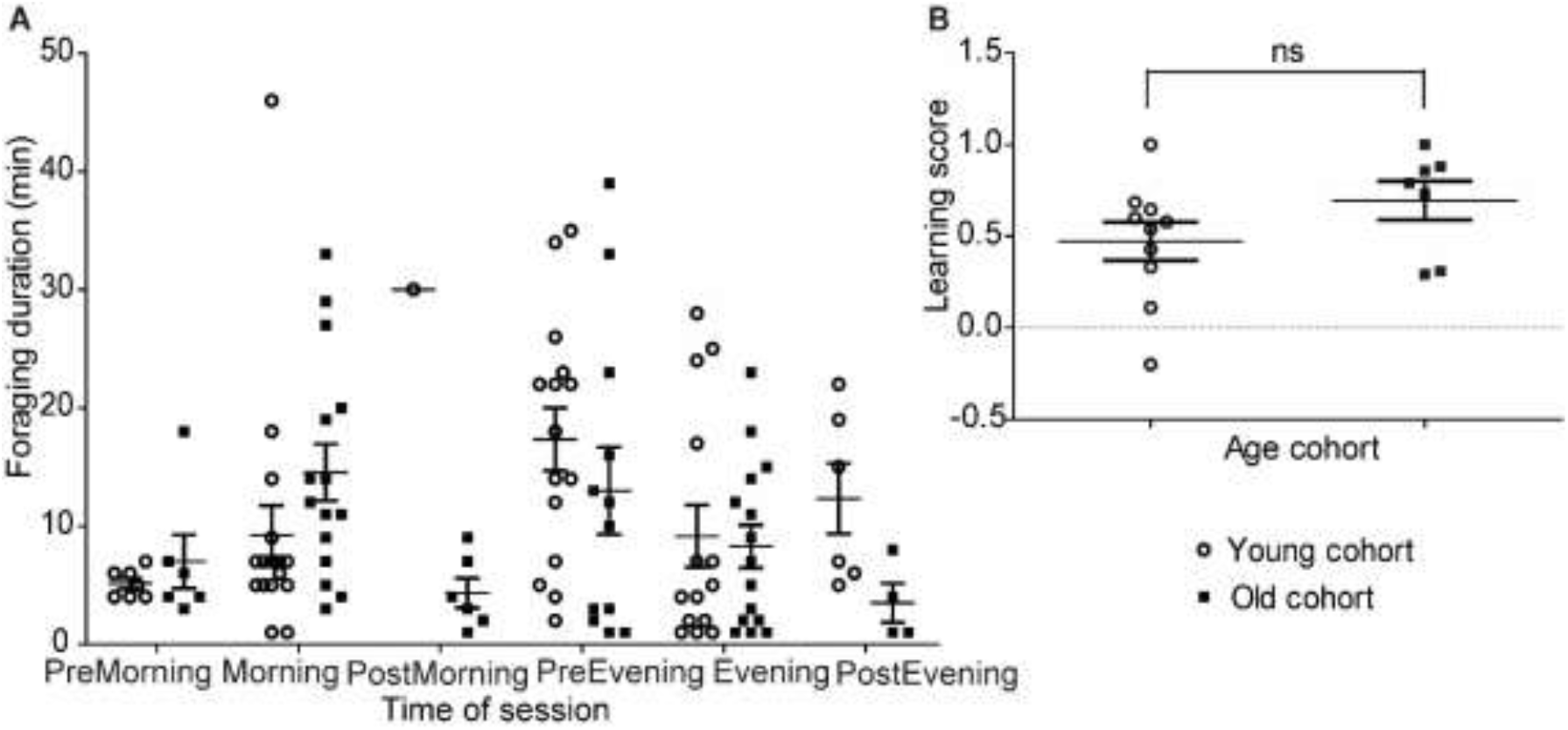
The influence of age cohort on learning performance and foraging activity. **A)** Foraging duration during the test day. The time sessions were pooled for the observations before the morning training (PreMorning), during the morning training (Morning), after the morning training (PostMorning), and before the evening training (PreEvening), during the evening training (Evening), and after the evening training (PostEvening). Both time of day and age cohort had a significant effect on foraging duration (2-way ANOVA, age *p* = 0.01, time *p* = 0.01, interaction *p* = 0.02). **B)** Learning scores during the time of the rewarding sessions. Performance during the morning and evening training sessions was pooled together. ns – not significant for Unpaired t-test, *p* = 0.16.

## Discussion

It is commonly assumed that time-memory is functionally significant. However, given that time-memory was studied for a very limited number of species, the generality of this capacity is not clear. We investigated the time-memory capacity in bumble bees which typically forage over a shorter time and distance than the better-studied honey bees. Our experiments lend credence to the hypothesis that bumble bees can learn the time of day and associate it with a reward and with external cues.

First, the colony in Exp. 2 showed the highest foraging activity during the rewarding hours and the lowest during the midday hours, and this pattern appeared different than in the control colony (*p*<0.06). Higher midday activity in the control compared to the trained colony was also suggested for the first experiment in which foraging activity was only assessed qualitatively. These findings showed that the time training affected the temporal organization of foraging activity in the focal colony (**Fig. 4**). These findings are consistent with studies with honey bees showing elevated foraging activity during rewarding times, and rest inside the hive for other times (Moore, 2001; Moore et al., 1989; Moore and Doherty, 2009; Wagner et al., 2013).

Second, the trained bees landed first on the color which was rewarding during this time of day during the training period (yellow during the morning, blue during the evening). The preference for the trained color was apparent already before the time of reward which is consistent with anticipation (**Fig. 5A**). Third, the total number of visits to the yellow and blue flowers differed with time of day with a clear preference for yellow during the morning sessions and for blue during the evening session. In this analysis too there was clear evidence for anticipation (**Fig. 3** and **Fig. 5**). Forth, in both experiments, bees showed a positive Learning Score during the rewarding sessions and the values were statistically different from zero in Exp. 2. These findings show that they learned to choose the color that was rewarding over the non-rewarding based on the time of day (**Fig. 3C** and **Fig. 5D**). Here too, there is good evidence for anticipation because their LS was positive already during the session just before the time of reward during both the morning and the evening. Studies on time-memory in honey bees show that trained bees typically arrive at the feeder before the time of reward with greater anticipation for the late-day food resources (Lehmann et al., 2011; Moore et al., 1989; Moore & Rankin, 1983). We showed that trained bumble bees anticipate the time of reward, which is characteristic of clock-regulated behavior. The Learning Score calculations in Exp. 2 are also consistent with more robust anticipation during the period preceding the evening compared to the morning training session (comparison of “PreMorning” and “PreEvening”, **Fig. 5D**).

It is worth mentioning that despite many differences (source colony, sample size, season, shape of artificial flowers, time of sunrise and sunset, light intensity, temperature throughout the day, humidity) between the two experiments, still, their results were overall similar and consistent with the hypothesis that bumble bees can associate time of day with sugar syrup reward and color. Our observations reveal individual variations among sister bees from the same colonies. To start exploring factors that may contribute to this individual variability, we compared the performance of bees of different body sizes and age cohorts. There was no sufficient size variability in our sample of foragers to allow for testing the influence of body size on learning performance. The low variability was probably also influenced by the association between body size and task performance in bumble bees with the larger bees more likely to forage (Chole et al., 2019; Free, 1955; Jandt and Dornhaus, 2009; Spaethe and Weidenmüller, 2002; Yerushalmi et al., 2006). Thus, additional studies with larger sample size and body size variability are needed in order to determine if the body size of the bee influences her performance in time-memory tasks. Our comparisons of bees from two age cohorts did not find an effect on the learning score, which is consistent with some previous studies. For example, age and body size did not affect associative odor learning tested by proboscis extension conditioning (Riveros and Gronenberg, 2009). Similarly, age and body size have no effect on ecologically relevant foraging performance and associative learning in bumble bees (Raine et al., 2006). We did find, however, an effect of age cohort on foraging duration with a significant interaction with time of day. These findings are consistent with the hypothesis that chronological age, experience, or both, influence the daily organization of foraging activity. Additional studies that are beyond the scope of the current work, are needed in order to understand the functional significance of these findings.

It is commonly accepted that time-memory is functionally significant because it allows animals to arrive at food patches at times of maximum reward while minimizing the time of risky foraging (reviewed in Mulder et al., 2013). The cost of arriving at the wrong time is specifically high for species such as honey bees with forage over long distances which also takes a lot of time. Thus, it is reasonable to assume that there is a selective advantage for precise timing visits to flowers for honey bees (reviewed in Bloch et al., 2017; Mulder et al., 2013). However, it was not clear whether similarly, strong selection pressures shaped time-memory in species typically foraging over shorter distances. Our evidence supporting the hypothesis that bumble bees associate time of day with a reward and other stimuli shows that efficient time-memory is not limited to species such as honey bees, which evolved sophisticated social foraging over long distances, and add to the relatively modest list of animals for which there is convincing empirical evidence for time-memory (Beling, 1929; Biebach et al., 1989; Chouhan et al., 2015; Harrison and Breed, 1987; Reebs, 1996; van der Zee et al., 2008). Similar comparative studies with species showing diverse life histories are necessary in order to understand the generality and adaptive significance of this complex clock-regulated behavior.

## Acknowledgments

This project was supported by the European Union’s Horizon 2020 research and innovation program (grant number: 765937-CINCHRON) and grants from US–Israel Binational Agricultural Research and Development Fund (BARD; IS-5077-18 R to Guy Bloch), and the US-Israel Binational Science Foundation (BSF; 2017188 to Guy Bloch).

## References

Allada R and Chung BY (2010) Circadian organization of behavior and physiology in Drosophila. Annu Rev Physiol. 72: 605–624.

Beekman M and Ratnieks FLW (2000) Long-range foraging by the honey-bee, Apis mellifera L. Funct Ecol. John Wiley & Sons, Ltd. 14: 490–496.

Beling I (1929) Uber das Zeitgedachtnis der bienen. J Comp Physiol.

Benirschke K (2004) Chronobiology: Biological Timekeeping. J Hered. Oxford University Press (OUP). 95: 91–92.

Biebach H, Gordijn M and Krebst JR (1989) Time-and-place learning by garden warblers, Sylvia borin. Anim Behav. 37: 353–360.

Bloch G (2009) Plasticity in the Circadian Clock and the Temporal Organization of Insect Societies. In: Organization of Insect Societies. Harvard University Press, pp. 402–432.

Bloch G, Bar-Shai N, Cytter Y, et al. (2017) Time is honey: Circadian clocks of bees and flowers and how their interactions may influence ecological communities. Phil Trans R Soc B. Royal Society Publishing. 372: 20160256.

Chole H, Woodard SH and Bloch G (2019) Body size variation in bees: regulation, mechanisms, and relationship to social organization. Current Opinion in Insect Science. Elsevier. 35: 77–87.

Chouhan NS, Wolf R, Helfrich-Förster C, et al. (2015) Flies Remember the Time of Day. Curr Biol. Cell Press. 25: 1619–1624.

Cueva Del Castillo R, Sanabria-Urb An S, Serrano-Meneses MA, et al. (2015) Trade-offs in the evolution of bumblebee colony and body size: a comparative analysis. Ecol Evol. 5: 3914–3926.

Dornhaus A and Chittka L (2004) Information flow and regulation of foraging activity in bumble bees (Bombus spp.). Apidologie. 35: 183–192.

Dornhaus A and Chittka L (2005) Bumble bees (Bombus terrestris) store both food and information in honeypots. Behav Ecol. 16: 661–666.

Dubowy C and Sehgal A (2017) Circadian rhythms and sleep in Drosophila melanogaster. Genetics. Genetics. 205: 1373–1397.

Free JB (1955) The division of labour within bumblebee Colonies. Insectes Sociaux. Springer. 2: 195–212.

Frisch K von (1967) The Dance Language and Orientation of Bees. Harvard University Press.

Harrison JM and Breed MD (1987) Temporal learning in the giant tropical ant, Paraponera clavata. Physiol Entomol. 12: 317–320.

Helm B, Visser ME, Schwartz W, et al. (2017) Two sides of a coin: ecological and chronobiological perspectives of timing in the wild. Phil Trans R Soc B. The Royal Society. 372: 20160246.

Jandt JM and Dornhaus A (2009) Spatial organization and division of labour in the bumblebee Bombus impatiens. Anim Behav. 77: 641–651.

Kleber E (1935) Hat das Zeitgedachnis der bienen biologische bedeutung? Zeitschrift fur vergleichende Physiologie. 22: 221–262.

Lehmann M, Gustav D and Galizia CG (2011) The early bee catches the flower -circadian rhythmicity influences learning performance in honey bees, Apis mellifera. Behav Ecol Sociobiol. Springer. 65: 205–215.

Manjunatha T, Dass H and Sharma SK (2008) Egg-laying rhythym in Drosophila melanogaster. J Genet. 87: 495–504.

Michener CD (1974) The Social Behavior of the Bees: A Comparative Study. Harvard University Press.

Moore D (2001) Mini-review Honey bee circadian clocks: behavioral control from individual workers to whole-colony rhythms. J Insect Physiol. 47: 843–857.

Moore D and Doherty P (2009) Acquisition of a time-memory in forager honey bees. J Comp Physiol A. 195: 741–751.

Moore D and Rankin MA (1983) Diurnal changes in the accuracy of the honey bee foraging rhythm. The Biological Bulletin. University of Chicago Press. 164: 471–482.

Moore D, Siegfried D, Wilson R, et al. (1989) The Influence of Time of Day on the Foraging Behavior of the Honeybee, Apis mellifera. J Biol Rhythms. 4: 305–325.

Mulder C, Gerkema MP and van der Zee EA (2013) Circadian clocks and memory: Time-place learning. Front Mol Neurosci. Frontiers. 6.

Pearce RF, Giuggioli L and Rands SA (2017) Bumblebees can discriminate between scent-marks deposited by conspecifics. Sci Rep. Nature Publishing Group. 7: 43872.

Raine NE, Ings TC, Ramos-Rodriguez O, et al. (2006) Intercolony variation in learning performance of a wild british bumblebee population (Hymenoptera: Apidae: Bombus terrestris audax). Entomol. E. Schweizerbart’sche Verlagsbuchhandlung. 28: 241–256.

Reebs SG (1996) Time-place learning in golden shiners (Pisces: Cyprinidae). Behav Processes. Elsevier B.V. 36: 253–262.

Renner MA and Nieh JC (2008) Bumble bee olfactory information flow and contact-based foraging activation. Insectes Sociaux. 55: 417–424.

Riveros AJ and Gronenberg W (2009) Olfactory learning and memory in the bumblebee Bombus occidentalis. Naturwissenschaften.Springer. 96: 851–856.

Saleh N, Scott AG, Bryning GP, et al. (2007) Distinguishing signals and cues: bumblebees use general footprints to generate adaptive behaviour at flowers and nest. Arthropod Plant Interact. Springer Science and Business Media LLC. 1: 119–127.

Saunders DS, Steel CGH, Vafopoulou X, et al. (2002) Insect Clocks. In: Insect Clocks. Elsevier, pp. 449–472.

Schmitt U (1990) Hydrocarbons in tarsal glands of Bombus terrestris. Experientia. 46.

Sharma VK (2003) Adaptive Significance of Circadian Clocks. Chronobiol. 20: 901–919.

Shpigler HY, Yaniv A, Gernat T, et al. (2022) The Influences of Illumination Regime on Egg-laying Rhythms of Honey Bee Queens. J Biol Rhythms. SAGE Publications Inc. 36226630.

Spaethe J and Weidenmüller A (2002) Size variation and foraging rate in bumblebees (Bombus terrestris). Insectes Sociaux. Birkhauser Verlag Basel. 49: 142–146.

Tataroglu O and Emery P (2014) Studying circadian rhythms in Drosophila melanogaster. Methods. Academic Press. 68: 140–150.

Time and date (2020a) Sunrise and sunset times in Jerusalem. Available at: https://www.timeanddate.com/sun/israel/jerusalem?month=9&year=2020&as_qdr=y15 (accessed 2 August 2022).

Time and date (2020b) Weather in Jerusalem. Available at: https://www.timeanddate.com/weather/israel/jerusalem/historic?month=9&year=2020 (accessed 3 August 2022).

Time and date (2021a) Sunrise and sunset times in Jerusalem. Available at: https://www.timeanddate.com/sun/israel/jerusalem?month=7&year=2021 (accessed 1 December 2022).

Time and date (2021b) Weather in Jerusalem. Available at: https://www.timeanddate.com/weather/israel/jerusalem/historic?month=7&year=2021 (accessed 1 December 2022).

van der Zee EA, Havekes R, Barf RP, et al. (2008) Circadian Time-Place Learning in Mice Depends on Cry Genes. Curr Biol. 18: 844–848.

von Buttel-Reepen H (1900) Sind die Bienen Reflexmaschinen? Experimentelle Beitrage zur Biologie der Honigbiene. Arthur Georgi. 20.

Wagner AE, van Nest BN, Hobbs CN, et al. (2013) Persistence, reticence and the management of multiple time memories by forager honey bees. J Exp Biol. 216: 1131–1141.

Wahl O (1932) Beitrag zur Frage der biologischen Bedeutung des Zeitgedächtnisses der Bienen. Springer-Verlag. 18: 709–717.

Walther-Hellwig K and Frankl R (2000) Foraging habitats and foraging distances of bumblebees, Bombus spp. (Hym., Apidae), in an agricultural landscape. J Appl Entomol. John Wiley & Sons, Ltd. 124: 299–306.

Wolf S and Moritz RFA (2008) Foraging distance in Bombus terrestris L. (Hymenoptera: Apidae). Apidologie. 39: 419–427.

Woodard SH, Lozier JD, Goulson D, et al. (2015) Molecular tools and bumble bees: revealing hidden details of ecology and evolution in a model system. Mol Ecol. John Wiley & Sons, Ltd. 24: 2916–2936.

Yerushalmi S and Green RM (2009) Evidence for the adaptive significance of circadian rhythms. Ecol. John Wiley & Sons, Ltd. 12: 970–981.

Yerushalmi S, Bodenhaimer S and Bloch G (2006) Developmentally determined attenuation in circadian rhythms links chronobiology to social organization in bees. J Exp Biol. The Company of Biologists. 209: 1044–1051.

