## Supplementary figures and images for "Bumble bees (*Bombus terrestris*) use time-memory to associate reward with color and time of day"

### Supplemental Figure 1

**Figure S1.** Experimental outline for Experiment 1.

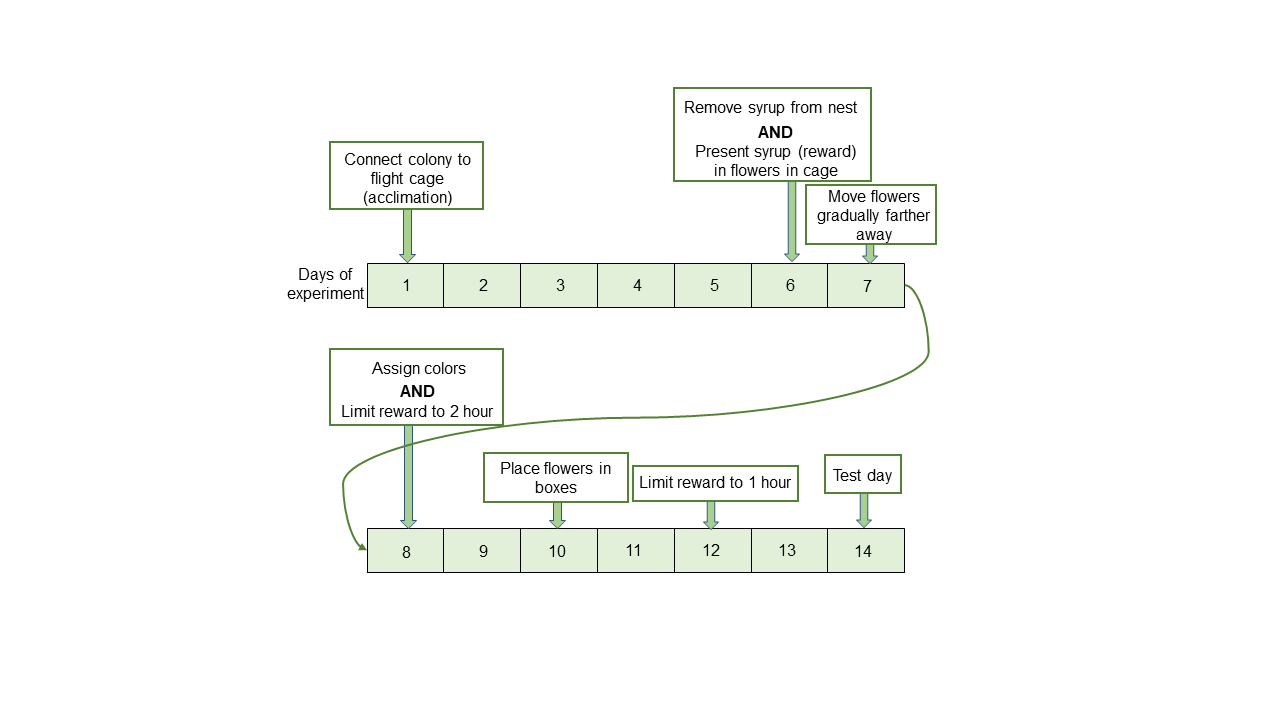

### Supplemental Figure 2

**Figure S2.** Experimental outline for Experiment 2.

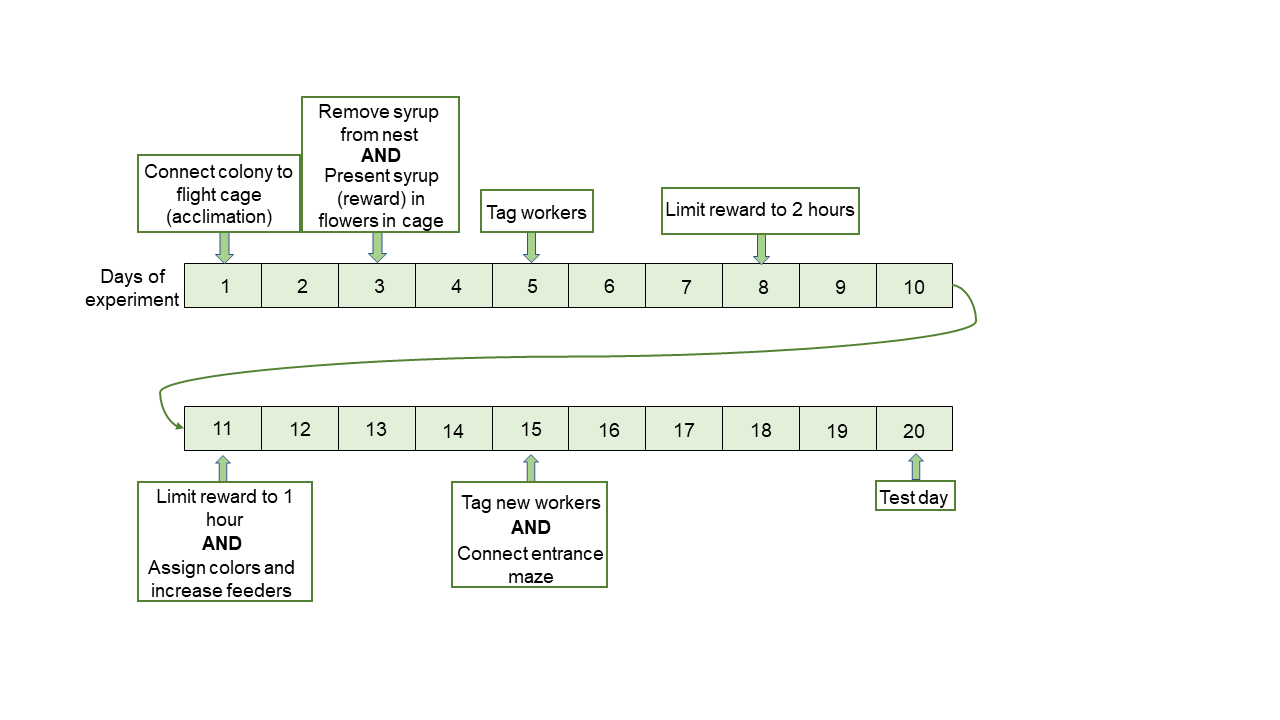
