## Supplemental Table 1 for "Bumble bees (*Bombus terrestris*) use time-memory to associate reward with color and time of day"

**Table S1 –** The total number of visits to ‘Yellow’ or ‘Blue’ flowers on the test day of Experiment 1. ‘Session’ refers to the period of the observations. ‘Yellow’ and ‘Blue’ columns show total visit counts to these colors. The ‘*p*.value’ gives the significance of a Chi-squared test assuming 50% to visit each one of the colors. ‘*p*.adj’ values are the p values after Bonferroni correction.

| Session | Yellow | Blue | *p*.value | *p*.adj |
| --- | --- | --- | --- | --- |
| PreMorning | 15 | 9 | 0.220 | 1 |
| Morning | 23 | 11 | 0.039 | 0.277 |
| PostMorning | 2 | 1 | 0.563 | 1 |
| Midday | 1 | 1 | 1 | 1 |
| PreEvening | 16 | 22 | 0.330 | 1 |
| Evening | 14 | 25 | 0.078 | 0.547 |
| PostEvening | 2 | 5 | 0.256 | 1 |
