## Supplemental Table 3 for "Bumble bees (*Bombus terrestris*) use time-memory to associate reward with color and time of day"

**Table S3 –** Training data for the second experiment, including all tagged foragers that visited the flowers. ‘Session count’ summarizes the number of observation sessions (morning or evening) in which the tagged bee visited the correct color at least once. ‘Old cohort’ and ‘Young cohort’ are the ID numbers of bees visiting flowers. Gray shaded IDs are trained bees identified during the test day.

| **Old cohort** | **Session count** | **Young cohort** | **Session count** |
| --- | --- | --- | --- |
| 8 | 9 | 12 | 2 |
| 11 | 6 | 18 | 2 |
| 14 | 1 | 42 | 1 |
| 17 | 1 | 52 | 1 |
| 22 | 17 | 54 | 2 |
| 28 | 14 | 58 | 6 |
| 35 | 15 | 64 | 1 |
| 37 | 19 | 65 | 1 |
| 38 | 3 | 73 | 2 |
| 42 | 1 | 79 | 4 |
| 46 | 8 | 81 | 3 |
| 51 | 1 | 83 | 8 |
| 53 | 12 | 87 | 4 |
| 58 | 12 | 89 | 5 |
| 60 | 10 | 96 | 1 |
| 72 | 1 | 97 | 2 |
| 74 | 15 |  |  |
| 79 | 2 |  |  |
| 80 | 1 |  |  |
| 84 | 2 |  |  |
| 86 | 15 |  |  |
