## Supplemental Table 2 for "Bumble bees (*Bombus terrestris*) use time-memory to associate reward with color and time of day"

**Table S2 –** Summary of the first choice of bees in the test day of the second experiment. Table details are as in Table S1.

| **Session** | **Yellow** | **Blue** | ***p*.value** | ***p*.adj** |
| --- | --- | --- | --- | --- |
| 06:30 | 9 | 0 | **0.002** | **0.032** |
| 07:00 | 10 | 4 | 0.108 | 1 |
| 07:30 | 17 | 0 | **3.74E-05** | **0.001** |
| 08:00 | 7 | 3 | 0.205 | 1 |
| 08:30 | 3 | 14 | **0.007** | 0.091 |
| 09:00 | 6 | 10 | 0.317 | 1 |
| 16:30 | 2 | 10 | **0.02** | 0.251 |
| 17:00 | 2 | 14 | **0.002** | **0.032** |
| 17:30 | 1 | 14 | **0.001** | **0.009** |
| 18:00 | 0 | 12 | **0.001** | **0.006** |
| 18:30 | 0 | 12 | **0.001** | **0.006** |
| 19:00 | 3 | 2 | 0.654 | 1 |
